# Effects of lithium on locomotor activity and circadian rhythm of honey bees

**DOI:** 10.1101/2023.05.17.541141

**Authors:** Babur Erdem, Okan Can Arslan, Sedat Sevin, Ayse Gul Gozen, Jose L. Agosto, Tugrul Giray, Hande Alemdar

## Abstract

The miticide effect of lithium on the honey bee (*Apis mellifera*) parasite *Varroa* has been discovered. *Varroa* mite is considered the principal threat to bee health and, as a result, to pollination and food security. In this study, we investigated the behavioral effects of lithium on honey bees, specifically locomotor activity (LMA) level and circadian patterns of LMA, such as rhythmicity, and periodicity. Any effects of lithium on activity may be important for bee health since timing and daylight are critical for bee foraging and bee’s use of the sun compass for navigation and communication. Both acute and chronic lithium treatments affected the LMA of honey bees. The effect varies depending on light and dark conditions. Chronic treatment with lithium disrupted the rhythmicity and altered the period of the circadian rhythm. While the circadian period was not affected by the light condition in the control group, lithium treatment lengthened the period in constant light condition. Lithium decreased total LMA in a constant light condition where typically activity is increased and not under the dark condition, both in acute and chronic treatments. However, mortality in the high-dose lithium treatment group is higher in the dark environment in the acute experiment. Lithium is also the first-line therapy for bipolar disorder. This disorder causes excessively elevated activity called mania and circadian rhythm abnormalities. The effects of lithium on reducing light-induced activity and the circadian rhythm of bees are reminiscent of its stabilizing effect on activity and circadian rhythms in bipolar disorder treatment.

## INTRODUCTION

Lithium is popularly known as the first-line treatment for bipolar disorder because of its mood-stabilizing activity and low toxicity. Lithium’s lethal effect on Varroa, a devastating parasite of honey bees, was surprisingly discovered by a research group while trying to find an iRNA-based treatment for Varroa control (Ziegelmann et al., 2018). Following the discovery, many studies focused on lithium’s residue amounts and toxicity on the *Varroa* and the honey bee at the colony level (Kolics et al., 2020; 2021a;b). Ziegelmann et al. fed bees 10 µl sucrose solution with at least 25 mM LiCl concentration and found that chronic feeding reduces their lifespan, but the 24- hour application has no effect (2018). Presern et al. also fed the bees with the same solution in the same amount. After 7 days, lithium concentrations were measured in the head, thorax, and abdomen as 6.50, 13.10, and 40.10 mg/kg, respectively. In the *ad libitum* application, bees were fed with the syrup for 7 days. Lithium concentrations were measured in the head, thorax, and abdomen as 39.25, 29.52, and 266.00 mg/kg, respectively (2020). Yet, there are very few studies on the behavioral effects of lithium applications on honey bees in the literature. One behavioral study investigated the lithium-induced malaise behavior of honey bees. In that study, lithium was applied via injection or feeding. The 0.1 or 1 mM LiCl was injected subcuticular in the thorax with 1 µl of the treatment solution. Each bee was fed 5 µl of a 1 M sucrose solution containing 0.1 or 1 mM LiCl. In this study, injection and ingestion of LiCl decreased walking and increased stillness. While injection increased the upside-down behavior, ingestion did not affect the behavior (Hurst et al., 2014). Only one study was conducted on the acute lithium salt effect on locomotor activity (LMA). We fed the bees with lithium formate (5μL), lithium lactate, or lithium citrate solutions. Bees fed with lithium citrate had higher LMA than lithium lactate (Sevin et al., 2022). However, the separate effects of citrate, formate, and lactate on LMA are unknown. Those first studies indicate that, beyond apiculture, perhaps we gain insights into lithium’s action as a drug from research on honey bees.

This study examined lithium’s effect on LMA and honey bees’ circadian rhythm. Our motivation stems from the moderating effect of lithium on bipolar disorder symptoms. Bipolar disorder is characterized by repetitive shifting between unusual mood levels as depression and mania. Irritable mood and abnormally and persistently increased activity or energy and decreased need for sleep are observed in manic episode (American Psychology Association, 2022). In addition to spikes of activity, circadian rhythm abnormalities also accompany bipolar disorder (McClung, 2007). Lithium is used as first-line therapy for bipolar disorder, and its stabilizing effect on activity is well known. Moreover, lithium has the ability to decelerate the excessively rapid circadian rhythms observed in individuals with bipolar disorder, as noted in previous studies (Atkinson et al., 1975; Kripke et al., 1978). However, the mechanism of action of lithium is only partially understood. This is partly due to the limited number of suitable animal models for bipolar disorder (Alda, 2015). Fruit fly (*Drosophila melanogaster*) is a powerful insect model in disease research. However, few studies use fruit flies to research lithium’s effects on locomotor activity (LMA), circadian rhythm, and behavior.

In flies, according to Dokucu et al., 20 and 30 mM LiCl significantly increase the period of the circadian rhythm to 24.02 h and 24.4 h, respectively, when compared to the control group, which has a period of 23.7 h. Also, lithium increases arrhythmicity, especially in doses higher than 5 mM, and it exceeds 50% at 60 mM application (2005). Another study on fruit flies indicates that the period of the circadian rhythm increases *Shaggy* (*Sgg*, insect orthologue of vertebrate *Gsk-3*) loss of function mutants (Martinek et al., 2001). It is known that the disruption of circadian rhythm is frequently observed in bipolar disease (Melo et al., 2017). Furthermore, circadian preference, a subjective preference for activities in the morning or evening, appears on the eveningness chronotype. Alteration in sleep latency and sleep/wake period reversal is observed in bipolar disorder (Giglio et al., 2010). Anomalies in the phase of circadian rhythms are modified with lithium treatment acting as an *Sgg* inhibitor (Escamilla & Zavala, 2008). In a study on longevity and locomotion, it was determined that the life expectancy of the fruit fly increased by *Sgg* inhibition after the application of 1–25 mM LiCl. In addition, it was observed that the treatment ameliorated the decrease in LMA due to aging (Castillo-Quan et al., 2016). In summary, lithium lengthens the period, increases longevity, and positively affects locomotion via *Sgg* inhibition in fruit flies (Jans et al., 2021).

Honey bees could be a possible insect model for studies on the effect of lithium on LMA, circadian rhythm, and behavior. Firstly, honey bees are sensitive to light regimes and circadian rhythms (see below). It is known that foraging activity strictly depends on the time of day when a flower provides nectar (Buttel-Reepen, 1915), resulting the maximizing the collected amount of it. Also, daylight is crucial for accurately navigating to and from nectar and pollen resources and communicating their location to nest mates using the sun-compass orientation (Bloch et al., 2017). Considering the central role of nectar and pollen foraging, orientation, and dance communication, individual activity and its daily regulation are vital in the long-term survival of the honey bee colony. The circadian rhythm of honey bees is susceptible to chemicals (Tackenberg et al., 2020; Chicas-Moiser et al., 2018) making this regulation an important research subject for bee health and accessible to mechanistic research on circadian rhythms. Honey bees have already provided insight into other complex behaviors. For example, cocaine affects the perception of reward in honey bees as it does in humans and withdrawal-like responses are also observed (Barron et al., 2009). Moreover, honey bees were successfully used as an ethanol-abuse model (Mixson et al., 2010). Dopamine, a biological amine common to humans and bees, has been found to affect learning and motivation in honey bees (Agarwal et al., 2011). The behavioral similarities between humans and honey bees are also mechanistically relevant, as shown recently that the socially unresponsive worker bees have similar gene expression patterns for the homologous versions of the autism-related genes in humans (Shpigler et al., 2019).

Based on the drug effects of lithium and other model animal studies, we hypothesize that the lithium treatment on bees may have more subtle effects than elevated mortality and alter LMA and circadian rhythms. We further hypothesize that the effectiveness of lithium treatment on LMA varies depending on light conditions, and long-term lithium treatment alters the circadian rhythm. To test our hypotheses, we altered the light conditions in short- and long-term experiments to examine LMA and circadian rhythm changes. Acute lithium treatment was applied to the honey bees, then, the bees were kept either in constant dark or light conditions for 24 hours in the short-term LMA experiments. We compared the LMA levels with different dosages of lithium for both light and dark conditions. In the long-term LMA experiment, honey bees were fed with lithium chronically for 15 days. This period was divided into 3 consecutive conditions: 12 hours of light and 12 hours of dark condition (12:12 LD), constant dark (DD), and constant light conditions (LL), respectively. Later, LMA activity levels, the honey bees’ rhythmicity of circadian rhythm, and the circadian period were analyzed statistically.

## RESULTS

### Acute effects of lithium on locomotor activity

In acute LMA trials, we used 3 doses of LiCl (low, medium, and high) and 1 dose of NaCl with a control group. The low, medium, and high doses of LiCl were 0.05 M, 0.15 M, and 0.45 M, respectively. The concentration of the NaCl applied in the acute LMA experiment was 0.45 M.

The LMA measurements did not follow the normal distribution (Shapiro-Wilk test, *p* < .05). A non-parametric Kruskal-Wallis test was used to compare the treatment groups for light and dark environment experiments.

First, we compared the LMA differences in the dark environment experiment. According to the Kruskal-Wallis test, there was no significant difference among the groups (*H* (4) = 9.11, *p* = .060). (Figure 1A).

**Figure 1.**
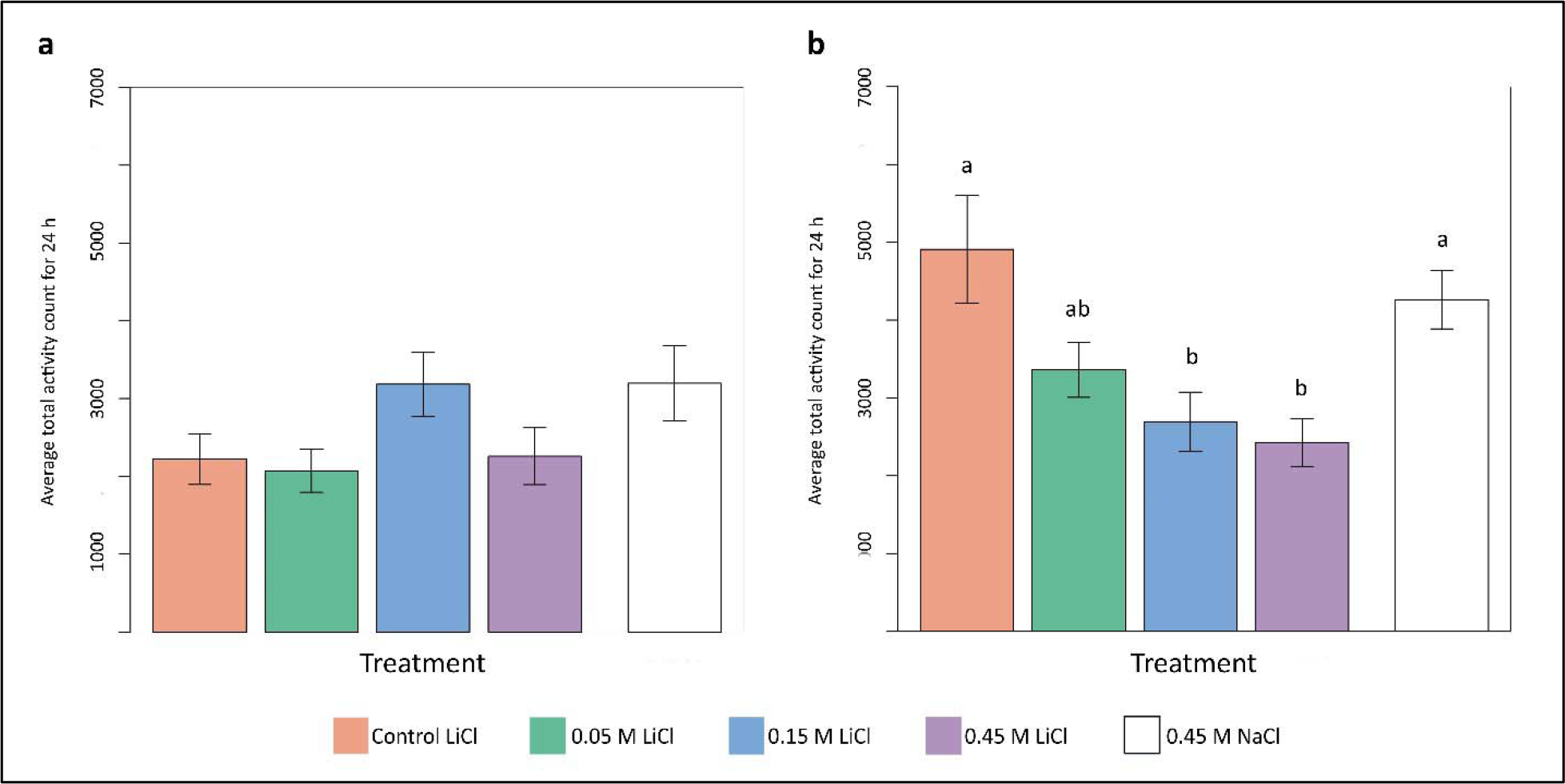
LMA comparisons of the acute experiments. There was no statistically significant found in both Kruskal-Wallis (p > .05) and regression analysis (p > .05) in the dark environment (a). However, according to the Kruskal-Wallis test, a statistical difference was found between the groups in the light environment (p < .001). Different letters on the bars indicate that groups differ according to a post hoc Dunn test (b). Also, regression analysis indicates a significant association between dosages of lithium in the light environment (b). Data are represented as the mean of total activity count and ± standard error.

Next, we compared the LMA of the groups in the light environment, and we found significant differences (Figure 1B) according to the Kruskal-Wallis test (H (4) = 16.86, *p* < .001). A post-hoc Dunn test indicated control group differed from medium (*p* = .004) and high dose groups (*p* < .001), similarly NaCl group differed from medium (*p* = .004) and high dose groups (*p* < .001), and there was no difference observed between control and NaCl groups (*p* = .486).

Then, we investigated the relationship between doses of LiCl (the NaCl group was excluded) and LMA using regression analysis. In dark environment experiments, there was not a significant association between dosages of lithium and activity (*R^2^* = .0001, *F* (1, 92) = .01, *p* = .916). However, the regression analysis indicated that there was a significant association between dosages of lithium and activity (*R^2^*= .1, *F* (1, 84) = 9.18, *p* = .003) in the light environment experiments. As a result, increasing the dosage decreased the activity in the light environment, but did not affect activity in the dark environment.

In addition, a chi-squared test was applied to find any difference across all groups, including both dark and light environment experiments regarding the mortality ratio in the groups. The test indicated a difference across the groups (χ*^2^* (9, 263) = 66.40, *p* < .001). As seen in Supplementary Table S1, the high-dose group in the dark environment had a higher death ratio and appeared as an outlier (Supplementary Figure S1, Supplementary Figure S2).

### Chronic effects of lithium on locomotor activity and circadian rhythm

In chronic LMA experiments, we compared 3 doses: low (1 mmol/kg), medium (5 mmol/kg), and high (10 mmol/kg), and a control group (no lithium administered).

We first analyzed the effect of lithium on survival. According to the survival analysis, the high-dose group had a lower survival rate (0.34) than other groups (control: 0.63, low: 0.56, medium: 0.63) on the 15th day. Difference between the groups was indicated by a log-rank test (χ2 = 7.9, *p* = 0.05) (Supplementary Figure S3), but according to pairwise comparisons, none of the lithium groups differs from the control (*p* > .05) (Supplementary Table S2).

Second, we compared the alteration of locomotor activities under the effect of different doses in different conditions. We used a repeated measure ANOVA test with eliminated random effects. Significant results were found for conditions (*F* (2, 34) = 8.54, *p* < .001), doses (*F* (3, 138) = 4.15, *p* = .008), and interaction effect (*F* (6, 138) = 2.69, *p* = .017). We applied a permutation test to our model because according to the Shapiro-Wilk test, the distribution was not normal (*p* < .05). The permutation test confirmed the repeated measures ANOVA test results for condition (B = 5000, *p* = .004), doses (B = 5000, *p* = .041), and interaction effect (B = 5000, *p* = .009). Then, we performed pairwise comparisons using Wilcoxon rank-sum exact test with the Holm adjustment method. We observed no difference (*p* > .05) among the low and medium-dose groups across LD, DD, and LL conditions. A slight decrease was observed in the DD condition compared to the LD condition for the high-dose group (*p* < .05). An apparent increase was found in the control group in the LL condition. The LMA measured for the control group in the LL condition was significantly higher than all other LMA measures of all conditions and all doses (*p* < .05) except the low dose LMA during the LL condition (*p* > .05) [Figure 2B] [Table 2] (Exact *p* values of the comparison for all pairs according to Wilcoxon test were given in supplement file, Supplementary Table S3).

**Figure 2.**
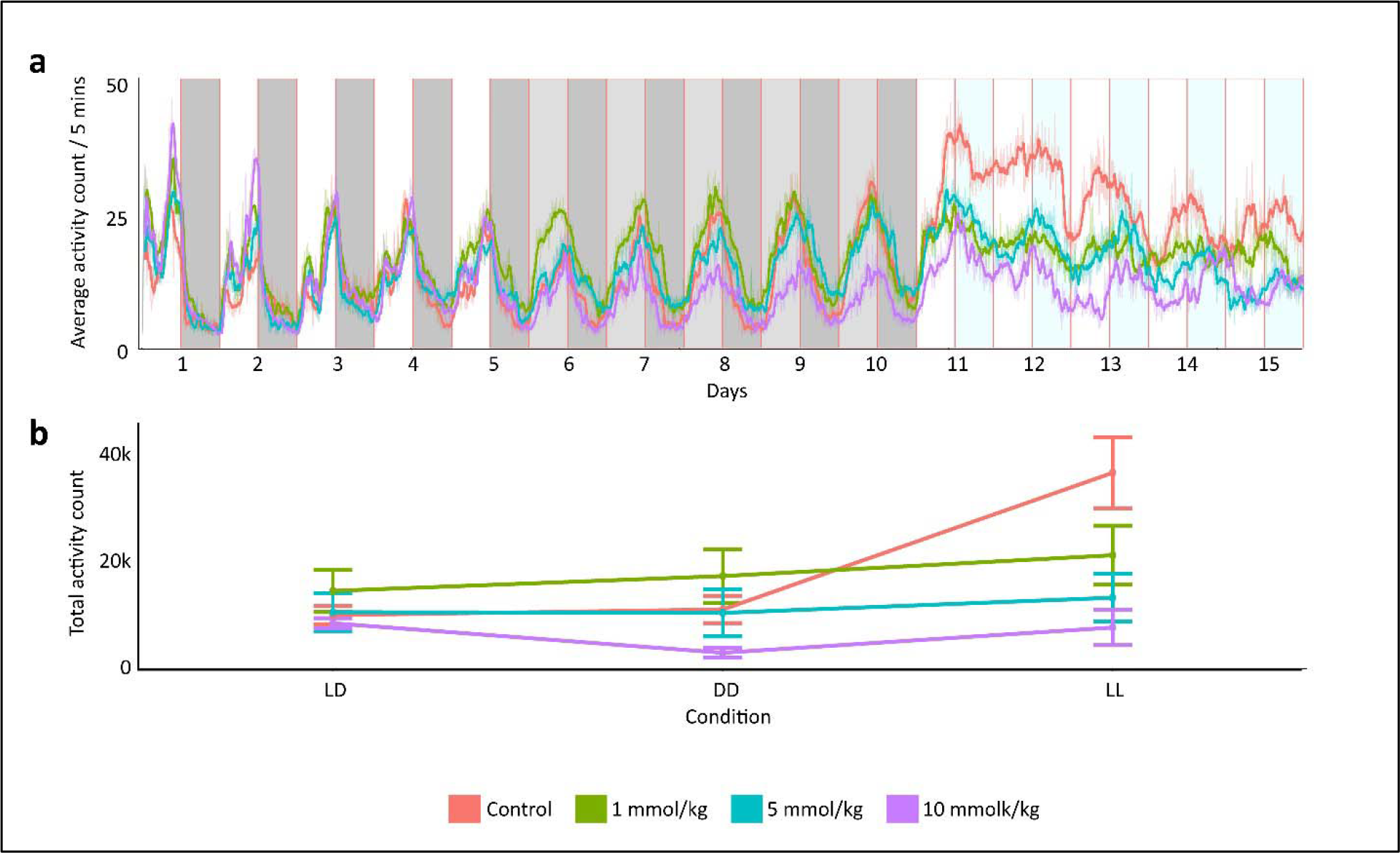
The activity graph represents changes in LMA during the chronic experiment. Dark or light grey shaded areas indicate darkness, white or light blue shaded areas indicate light periods, and the areas divide the plots into 12-hour intervals. A moving average filter was applied; it took 12 data points at a time as input. Thus, it produced an average activity of 60 min for a single output point (a). Total activity count in conditions as LD, DD, and LL, each lasting 5 days, was used in the pairwise Wilcoxon rank-sum exact test with the Holm adjustment method. There was no difference observed (p > .05) across conditions for low and medium doses. A slight decrease was observed in the DD condition of the high dose (p < .05) compared to the LD condition of the high dose. An apparent increase was found in the control group at the LL condition, which is significantly different from all data points of all conditions and all doses (p < .05) except the data point of low dose at the LL condition (p > .05). Data are represented as the mean of total activity count and ± standard error (b).

**Table 1.** Descriptive statistics for acute LMA experiments.

|  |  | n | Mean | Std.Dev. | Median | Std.Err. |
| --- | --- | --- | --- | --- | --- | --- |
| Dark<br>Environment<br>Experiments | Control | 24 | 2220.46 | 1596.29 | 1913.00 | 325.84 |
|  | 0.05M LiCl | 23 | 2070.65 | 1322.75 | 1721.00 | 275.81 |
|  | 0.15M LiCl | 22 | 3184.46 | 1932.33 | 2422.50 | 411.97 |
|  | 0.45M LiCl | 25 | 2259.08 | 1829.77 | 1748.00 | 365.95 |
|  | 0.45M NaCl | 24 | 3196.58 | 2364.28 | 2480.00 | 482.61 |
| Light<br>Environment<br>Experiment | Control | 22 | 4908.55 | 3246.16 | 3827.50 | 692.08 |
|  | 0.05M LiCl | 24 | 3360.38 | 1739.24 | 3134.50 | 355.02 |
|  | 0.15M LiCl | 20 | 2695.30 | 1694.88 | 2509.50 | 378.99 |
|  | 0.45M LiCl | 20 | 2426.75 | 1373.42 | 2477.00 | 307.11 |
|  | 0.45M NaCl | 24 | 4259.17 | 1824.97 | 3728.00 | 372.52 |

**Table 2.** Descriptive statistics for activity counts in chronic LMA experiments.

|  | Condition | n | Mean | St.Dev | Median |
| --- | --- | --- | --- | --- | --- |
| Control | LD | 20 | 12445.80 | 6994.88 | 11653.00 |
|  | DD | 20 | 13415.50 | 10570.17 | 11588.50 |
|  | LL | 20 | 37158.90 | 27481.49 | 25793.00 |
| 1 mmol/kg | DD | 18 | 19215.06 | 19739.43 | 9329.50 |
|  | LD | 18 | 16655.72 | 15558.48 | 12473.50 |
|  | LL | 18 | 22845.72 | 21835.67 | 10716.00 |
| 5 mmol/kg | LD | 20 | 12952.80 | 14754.55 | 9651.50 |
|  | DD | 20 | 12826.55 | 18271.93 | 7354.50 |
|  | LL | 20 | 15472.85 | 18533.57 | 9632.50 |
| 10 mmol/kg | LD | 11 | 11028.18 | 2903.24 | 11207.00 |
|  | DD | 11 | 5893.27 | 2844.56 | 5100.00 |
|  | LL | 11 | 10289.27 | 10167.60 | 5496.00 |

Third, we compared the rhythmicity of circadian rhythm ratios for each condition according to the Lomb–Scargle periodogram analysis result. We found the ratios of rhythmic individuals in the LD condition for control, low, medium, and high dose groups to be 1, 0.97, 0.90, and 1, respectively. In the DD condition, 0.96, 0.96, 0.93, and 0.75. In the LL condition, 0.85, 0.83, 0.70, and 0.73, respectively (Table 3, Figure 3A). Using the logistic regression model, we inspected the association between rhythmicity and lithium treatment in each condition. We found a significant association in the DD condition (χ*2* (3, 93) = 5.21, p = .023) but not in LD (χ*2* (3, 110) = 0.001, *p* = .974) and LL (χ*2* (3, 67) = 1.20, *p* = .273). Thus, rhythmicity significantly decreased in DD condition. In addition, although not statistically significant, a noticeable decrease was observed in the LL condition. (Table 3).

**Figure 3.**
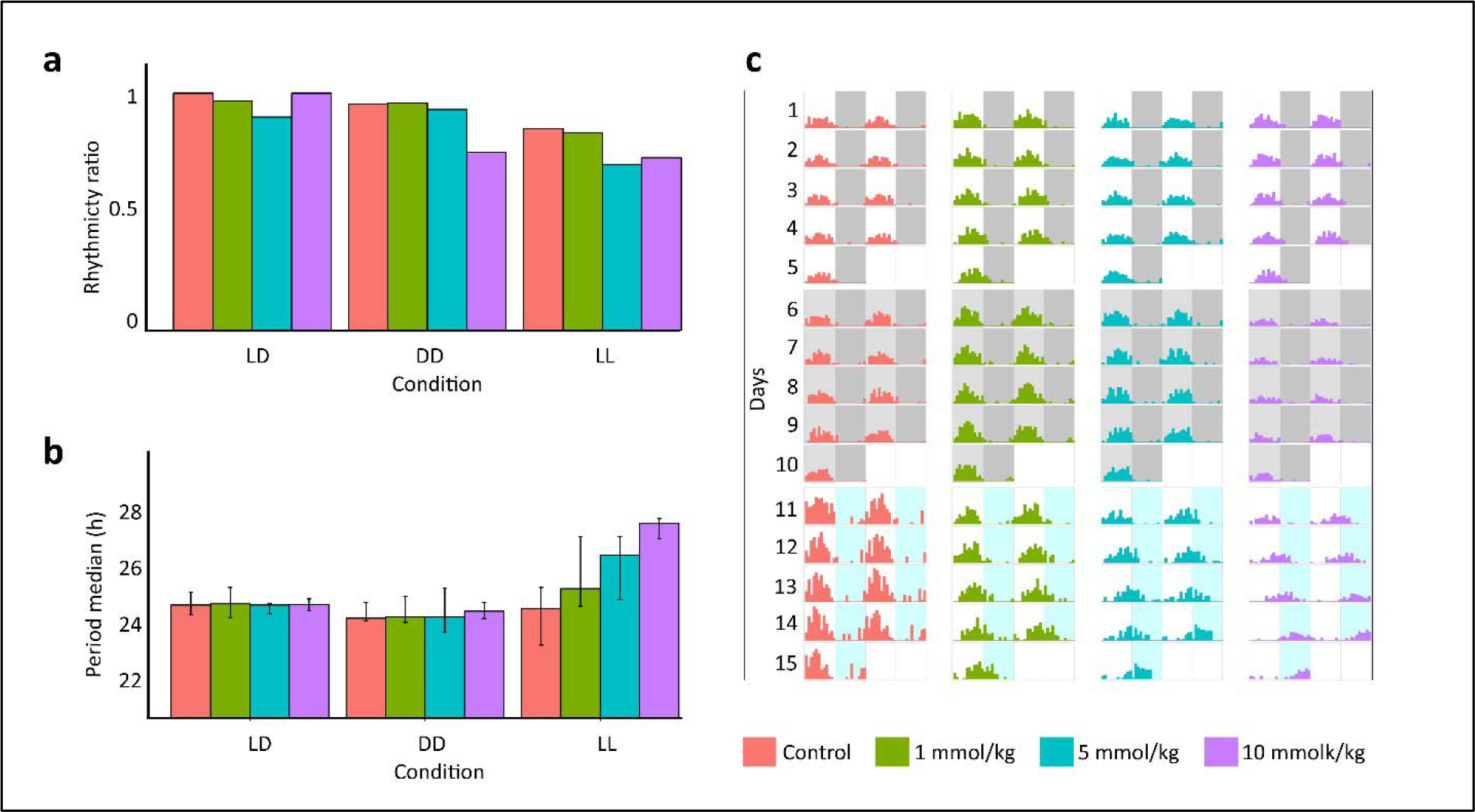
Rhythmicity significantly decreased in the DD condition according to the logistic regression model (p < .001), also although not statistically significant, a noticeable decrease was observed in the LL condition (a). According to the Spearman correlation, a significant relationship between periodicity and lithium treatment was observed only for the LL condition (p < .001), the medians are represented with the interquartile ranges (b). Double-plotted actograms of representative individuals were generated using the sine wave function; *y(t) = Asin(2*π*f +* φ*)+D*. Amplitude, center amplitude, and frequency variables were tuned according to the LMA averages and lengths of the circadian period of each dose group in each condition, and arrhythmicities were added as noise. Dark or light grey shaded areas indicate darkness, white or light blue shaded areas indicate light periods, and the areas divide the plots into 12-hour intervals (c).

**Table 3.**
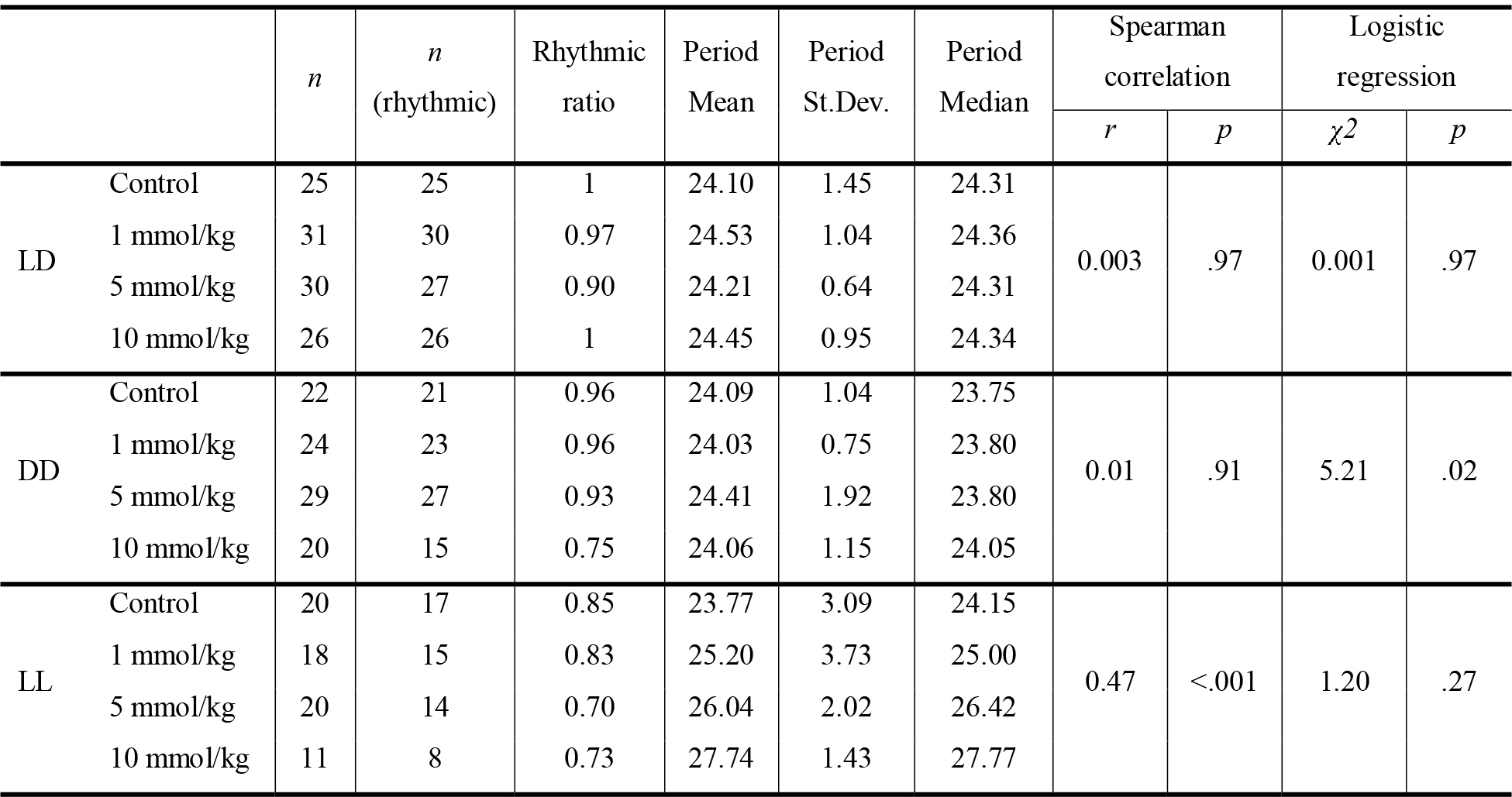
Descriptive statistics for chronic LMA experiment on rhythmicity and periodicity, results of the Spearman correlation test for periodicity, and logistic regression test for rhythmicity.

|  |  | *n* | *n*<br>(rhythmic) | Rhythmic<br>ratio | Period<br>Mean | Period<br>St.Dev. | Period<br>Median | Spearman<br>correlation |  | Logistic<br>regression |  |
| --- | --- | --- | --- | --- | --- | --- | --- | --- | --- | --- | --- |
| | | | | | | | | *r* | *p* | $\chi^2$ | *p* |
| LD | Control | 25 | 25 | 1 | 24.10 | 1.45 | 24.31 | 0.003 | .97 | 0.001 | .97 |
|  | 1 mmol/kg | 31 | 30 | 0.97 | 24.53 | 1.04 | 24.36 |  |  |  |  |
|  | 5 mmol/kg | 30 | 27 | 0.90 | 24.21 | 0.64 | 24.31 |  |  |  |  |
|  | 10 mmol/kg | 26 | 26 | 1 | 24.45 | 0.95 | 24.34 |  |  |  |  |
| DD | Control | 22 | 21 | 0.96 | 24.09 | 1.04 | 23.75 | 0.01 | .91 | 5.21 | .02 |
|  | 1 mmol/kg | 24 | 23 | 0.96 | 24.03 | 0.75 | 23.80 |  |  |  |  |
|  | 5 mmol/kg | 29 | 27 | 0.93 | 24.41 | 1.92 | 23.80 |  |  |  |  |
|  | 10 mmol/kg | 20 | 15 | 0.75 | 24.06 | 1.15 | 24.05 |  |  |  |  |
| LL | Control | 20 | 17 | 0.85 | 23.77 | 3.09 | 24.15 | 0.47 | <.001 | 1.20 | .27 |
|  | 1 mmol/kg | 18 | 15 | 0.83 | 25.20 | 3.73 | 25.00 |  |  |  |  |
|  | 5 mmol/kg | 20 | 14 | 0.70 | 26.04 | 2.02 | 26.42 |  |  |  |  |
|  | 10 mmol/kg | 11 | 8 | 0.73 | 27.74 | 1.43 | 27.77 |  |  |  |  |

Lastly, we applied a Lomb–Scargle periodogram analysis to determine the statistics of rhythmic individuals. According to periodogram analysis, we determined the median of the periods of the circadian rhythm with an interquartile range (Q1 – Q3) in hours. In the LD condition, the periods were 24.31 (23.90 – 24.84) for control, 24.36 (23.76 – 25.05) for low dose, 24.31 (23.95 – 24.39) for medium-dose, and 24.34 (24.08 – 24.57) for high dose bees. In the DD condition, the periods were 23.75 (23.65 – 24.41) for control, 23.80 (23.58 – 24.68) for low dose, 23.80 (23.17 – 25.03) for medium dose, 24.05 (23.74 – 24.41) for high dose bees. In the LL condition, the periods were 24.15 (22.62 – 25.05) for control, 25.00 (24.26 – 27.20) for low dose, 26.42 (24.56 – 27.20) for medium dose, and 27.77 (27.12- 27.96) for high dose bees (Figure 3B). Then, we determined the relationship between the periodicity and lithium treatment in each condition. We used a non-parametric Spearman correlation test because the normal distribution was not met according to the Shapiro-Wilk test (*p* < .05). There was no correlation in LD (*r* (106) = .003, *p* = .974), and DD (*r* (84) = .01, *p* = .908) conditions. However, a positive correlation was found in the LL condition (*r* (58) = .47, *p* < .001). This shows lithium affected the periodicity under constant light condition (Table 3, Figure 3B). Additionally, we compared the circadian period length of each dose across conditions. According to the Kruskal-Wallis test, the light condition was not effective on the length of the circadian period for the control (*H* (2) = 0.98, *p* = .61) and low dose group (*H* (2) = 4.18, *p* = .12). Effect of the light condition on circadian period length appeared in medium (*H* (2) = 11.40, *p* < .001) and high dose groups (*H* (2) = 19.70, *p* < .001). Post-hoc Dunn test indicated the length of the circadian period in LL condition differed from LD and DD conditions for both medium (*p* = .002 and *p* < .001) and high dose (*p* < .001 and *p* < .001) groups. Thus, period length is only increased by constant light condition for medium and high doses (Supplementary Figure S4).

## DISCUSSION

The principal finding of this study is that lithium alters the locomotor activity and circadian rhythms of honey bees. Acute lithium treatment decreased light-induced higher activity in honey bees, yet did not affect activity observed under no light conditions. Similarly, in the chronic experiment, lithium decreased the elevated locomotor activity in constant light condition. Furthermore, in addition to the locomotor activity, we also examined the effect of lithium on rhythmicity and the circadian period in the chronic experiment. The results indicated that lithium treatment disrupted rhythmicity by significantly lowering the rhythmicity ratio in the DD condition, and an apparent decrease was observed in the LL condition, although not statistically significant. The circadian period was not affected by different light conditions for bees in the control group, yet it lengthened with lithium treatment under the constant light condition.

While the discovery of lithium’s lethal effect on *Varroa* mites is significant for bee health (Ziegelmann et al., 2018), chronic exposure to lithium reduces bee lifespan, and high concentrations are toxic (Kolics et al., 2020; 2021a;b; Presern et al., 2020). Therefore, a more comprehensive understanding of the effects of lithium on honey bees is necessary before it can be considered a conventional acaricide. The current study’s results on lithium’s effect on locomotor activity and circadian rhythm are critical because locomotor activity and circadian rhythm play a crucial role in regulating bee foraging activity and navigation ability (Bloch et al., 2017). Our observation is important not only for understanding lithium’s effects on bees’ health but also for basic neuroscience research. It is notable to consider honey bees as a potential non-mammalian model for bipolar disorder, given the similarities in activity observed in bipolar disorder patients and animal models.

We propose using constant light to trigger mania-like behavior since exposure to light leads to higher activity (Spangler, 1973), as also observed in our study. Different methods were used to increase the activity to develop a mania model. The mania-like activity was triggered by paradoxical sleep deprivation (Armani et al., 2012), drugs (Smith, 1980), and mutations in the *Clk* gene important for circadian rhythms (Roybal et al. 2007). Lithium decreased the elevated activity in all three studies. In our study, similar to previous studies, lithium treatment reduced elevated activity under acute or chronic light conditions compared to control bees. In contrast, control and lithium-treated bees showed similar, lower locomotor activity under dark conditions. That is consistent also with lithium maintenance trials in patients with bipolar disorder: lithium is predominantly effective against mania instead of depression (Tondo et al., 2019; Bowden et al., 2003; Calabrese et al., 2003).

Aside from its effects on locomotor activity levels, lithium also impacts the strength of circadian rhythms and the length of the circadian period. Previous studies by Dokucu et al. (2005) and Smietanko & Engelmann (1989) have shown that lower rhythmicity was observed in fruit flies and house flies (*Musca domestica*), respectively, under lithium treatment in DD or low light conditions. In our study, we also observed the disruptive effect of lithium on rhythmicity. The possible molecular mechanism underlying this observation is attributed to lithium inhibition of the activity of vertebrate *Gsk-3* ortholog *Sgg* which is related to the regulation of signal transduction, xenobiotic stress resistance, and neuronal health (Jans et al., 2021). It is known to suppress arrhythmicity (Stoleru et al., 2007).

Chronic research on diurnal vertebrates such as goldfish (*Carassius auratus*), and squirrel monkeys (*Saimiri sciureus*), have found lengthening of the circadian period with lithium treatment, and these were measured in LL conditions (Kavaliers, 1981; Welsh & Moore-Ede, 1990). In our study, we observed an increase in the circadian period only in LL conditions, while in other insects under lithium treatment, the period increased in DD condition. Lithium has been shown to affect the circadian period length in various organisms, including nocturnal mammals such as mice and hamsters (*Mesocricetus auratus*), as well as insects such as cockroaches (*Leucophaea maderae*), fruit flies, and house flies, under DD or weak red light conditions (Dokucu et al., 2005; Hofmann et al., 1978; Iwahana et al., 2004; Padiath et al., 2004; Smietanko & Engelmann, 1989). Notably, we did not observe a statistically significant increase in the period in the LL condition (24.15 h) vs. the DD condition (23.8 h) in the control group, although lengthening of the circadian period under constant light in insects is a known general rule (Aschoff, 1979). These results suggest that honey bees may differ from other insects in their circadian period regulation. Honey bees carry the mammalian-type *Cryptochrome* gene *Cry2* but lack the *Cry1* and *Tim1* genes involved in circadian rhythm regulation in other insects (Beer & Bloch, 2020). *Cry1* loss-of-activity mutant fruit flies did not show period lengthening in LL conditions (Emery et al., 2000), indicating the absence of *Cry1* in honey bees may be the reason for not observing period lengthening in the control group under the constant light condition. The inactivation of *Sgg* causes circadian period lengthening in DD conditions (Martinek et al., 2001). Also, *Sgg* has a regulatory role on the *Cry1* and *Tim1* genes in fruit flies (Stoleru et al., 2007). The absence of these interactions in honey bees may be why we could not observe an increase in the circadian period in DD with lithium treatment. However, instead of light-sensitive *Cry1*, vertebrate-like opsin called pteropsin is predominantly found in honeybees (Velarde et al., 2005). Possible interactions between SGG and pteropsin may reveal the cause of the circadian period lengthening with lithium treatment under LL condition in bees, as observed in diurnal vertebrates.

Future research on honey bees should consider their unique circadian rhythm ontogeny, driven by age and social environment. In honey bees, the rhythmicity development depends on the worker bees’ roles in the colony. Young nurse bees do not display circadian rhythms, while older forager bees exhibit robust circadian oscillations in the expression of circadian-related genes such *Per*, *Cry2*, *Tim2*, and *Clk* (Bloch, 2010).

In brief, we evaluated the potential impact of lithium on the bees’ circadian rhythm and locomotor activity. We emphasize the need for careful consideration of the implications of using lithium for *Varroa* control to ensure the long-term sustainability of honey bee populations and their crucial role in global food security. Furthermore, we suggest that honey bees should be considered experimental animal models for psychological research due to their similarities in gene patterns and behavior to humans (Barron et al., 2008; Mixson et al., 2010; Agarwal et al., 2011; Shpigler et al. 2019). Overall, our research on lithium’s impact on honey bees has significant implications for honey bee health and potential use as an experimental model for neurobiological research.

## MATERIAL AND METHOD

### Sampling

Anatolian honey bee (*Apis mellifera anatoliaca*) hives in the apiary of Ankara University’s Veterinary Faculty sources were used in this study. Hive entrances were blocked with plastic wire meshes. Returning forager bees that could not enter the hive congregated on the wire meshes and were collected into small containers (Scheiner et al., 2013). Samples were from at least three hives and pooled for each analysis to mitigate the colony effect.

### Acute experiments

Bees were fed 10 μl of sucrose solution (50% w/v) containing 0.05, 0.15, and 0.45 M LiCl for treatments and the control group. In addition, we used the 0.45 M NaCl treatment as a control for the hyper-osmotic stress that may be caused by 0.45 M salt solution. The LMA analysis was performed as described in Giannoni-Guzman et al. (2014). After treatments, honey bees of experimental groups were individually transferred into perforated 15 ml falcon tubes. A piece of fondant sugar was placed into the cap of each tube as food and covered with two layers of cheesecloth to prevent the sticking of bees to the fondant sugar. The LMA monitoring device produced by Trikinetics Inc (TriKinetics Inc, Waltham, MA USA) was used in our study. We used 4 modules with 24 holes to hold a single 15 ml falcon tube. The three infrared sensors around each hole gave a positive signal when the bee in the Falcon tube moved through the section with the beams. An environmental monitor was also attached to the system, constantly measuring and recording temperature, light, and humidity levels. The LMA monitoring modules were kept in an incubator at 33 °C and 65% humidity.

The LMAs of bees in treatment and control groups were measured for 24 hours in two experiments, one in constant darkness and another in a constant light environment.

### Chronic experiments

The chronic LMA experiment was performed according to Tackenberg et al. (2020). Collected forager honey bees were put into 15 ml falcon tubes individually. A piece of custom-made bee candy was placed into the cap of each falcon tube and covered with two sheets of cheesecloth. Bee candy consisted of 10 parts of honey, 54 parts of powdered sugar, and 2 parts of distilled water in grams; this recipe slightly differed from that of Tackenberg et al. (2020). Designated amounts of LiCl were dissolved in the water fraction of treatment groups (1, 5, and 10 mmol/kg of LiCl). The chronic LMA experiment was performed in the same incubator (33 °C, 65% humidity) for 15 days with 5 days in 12-hour light and dark cycle (light: dark, LD) followed by 5 days in constant darkness (dark: dark, DD) and the final 5 days under continuous illumination (light: light, LL).

### Statistics

All statistical analyses were conducted in RStudio with the R 4.3 version (R Core Team, 2020). The normality of the sample distribution was checked with the Shapiro-Wilk test.

In acute experiments, in the case of the distributions that did not follow the normal distribution, the activity differences across the groups were compared by the non-parametric Kruskal-Wallis test followed by a post-hoc Dunn test. The relationship between the dosages and LMA was examined in a regression analysis. We checked group differences in mortality via Chi-Square analysis.

In chronic treatment experiments, a survival analysis was done by log-rank test to compare mortality differences. Alteration of activities under the effect of different lithium doses in different light exposure conditions was examined via repeated measure ANOVA tests with eliminated random effects. Then, pairwise comparisons were applied using Wilcoxon rank-sum exact test with the Holm adjustment method. Also, a permutation test was used to verify the repeated measure ANOVA test results. Circadian rhythm analysis determining the periodicity and rhythmicity was conducted with a set of R packages called “Rethomics”, available at https://rethomics.github.io (Geissman et al., 2019). Lomb–Scargle periodogram analysis was used to determine the rhythmic individuals and lengths of the circadian periods. Then, the logistic regression model was used to determine the association between rhythmicity and lithium treatment. The Spearman correlation test was used to determine the association between the circadian period and lithium treatment for each condition.

## DATA AVAILABILITY

The datasets generated during and/or analysed during the current study are available in the Zenodo repository, https://doi.org/10.5281/zenodo.7939046

## ACKNOWLEDGEMENTS

This work was supported by The Scientific and Technological Research Council of Türkiye [TÜBİTAK grant number: 122E014]; and the Middle East Technical University Research Fund [BAP grant numbers: HDESP-312-2021-10868; GAP-312-2022-10845].

## AUTHOR CONTRIBUTIONS

BE, TG, and HA designed research. BE, OCA, and SS performed experiments. BE analyzed data. BE, TG, HA, and JLA interpret the results. BE, TG, HA, and AGG wrote and revised the manuscript. All authors reviewed the manuscript.

## SUPPLEMENTS

### Acute experiments

**Table S1.**
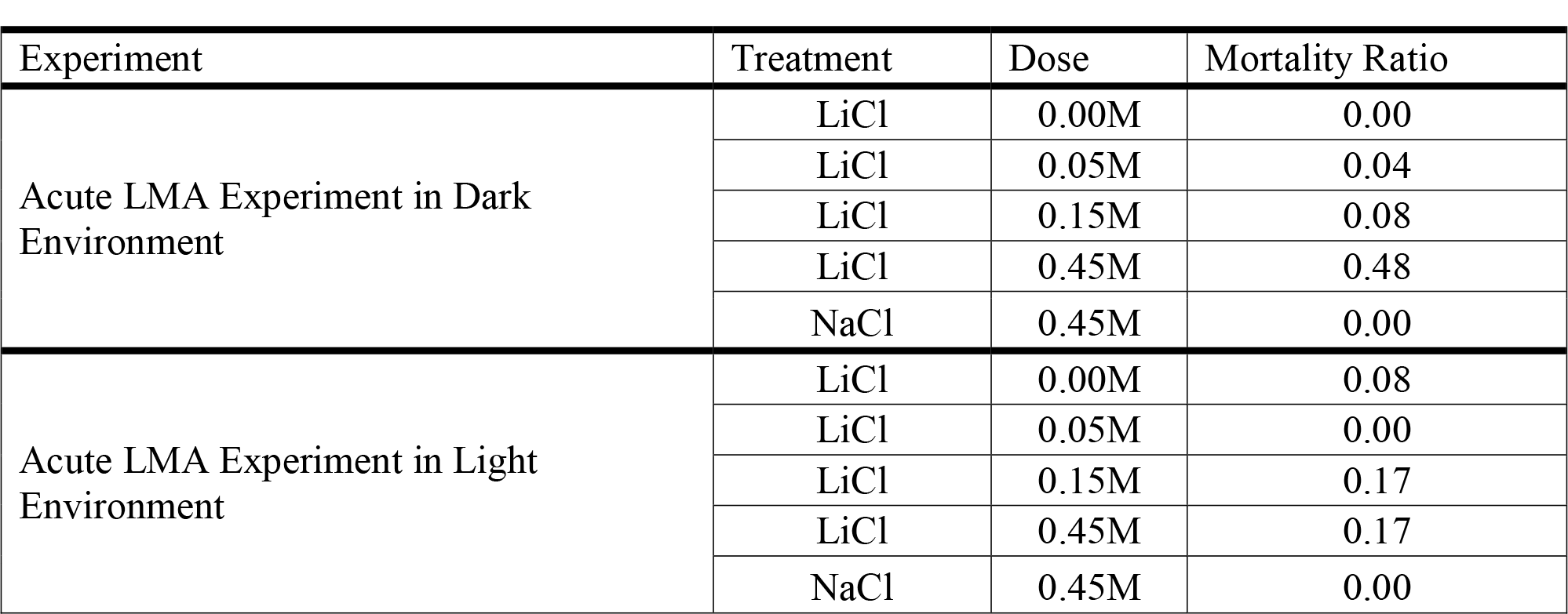
Mortality ratios of the groups acute LMA experiments in the dark and light environment.

**Figure S1.**
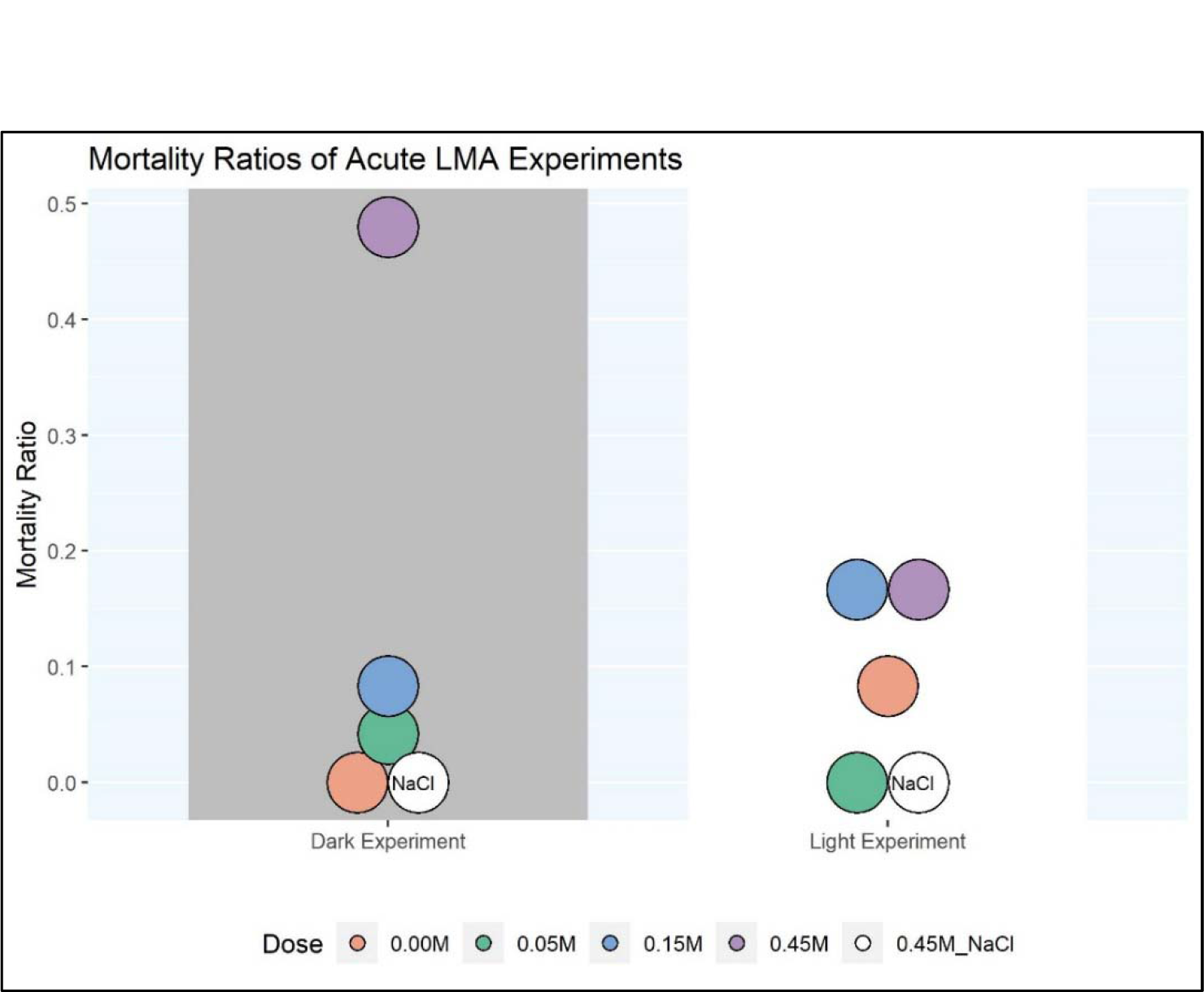
Mortality ratios of the groups acute LMA experiments in the dark and light environment.

**Figure S2.**
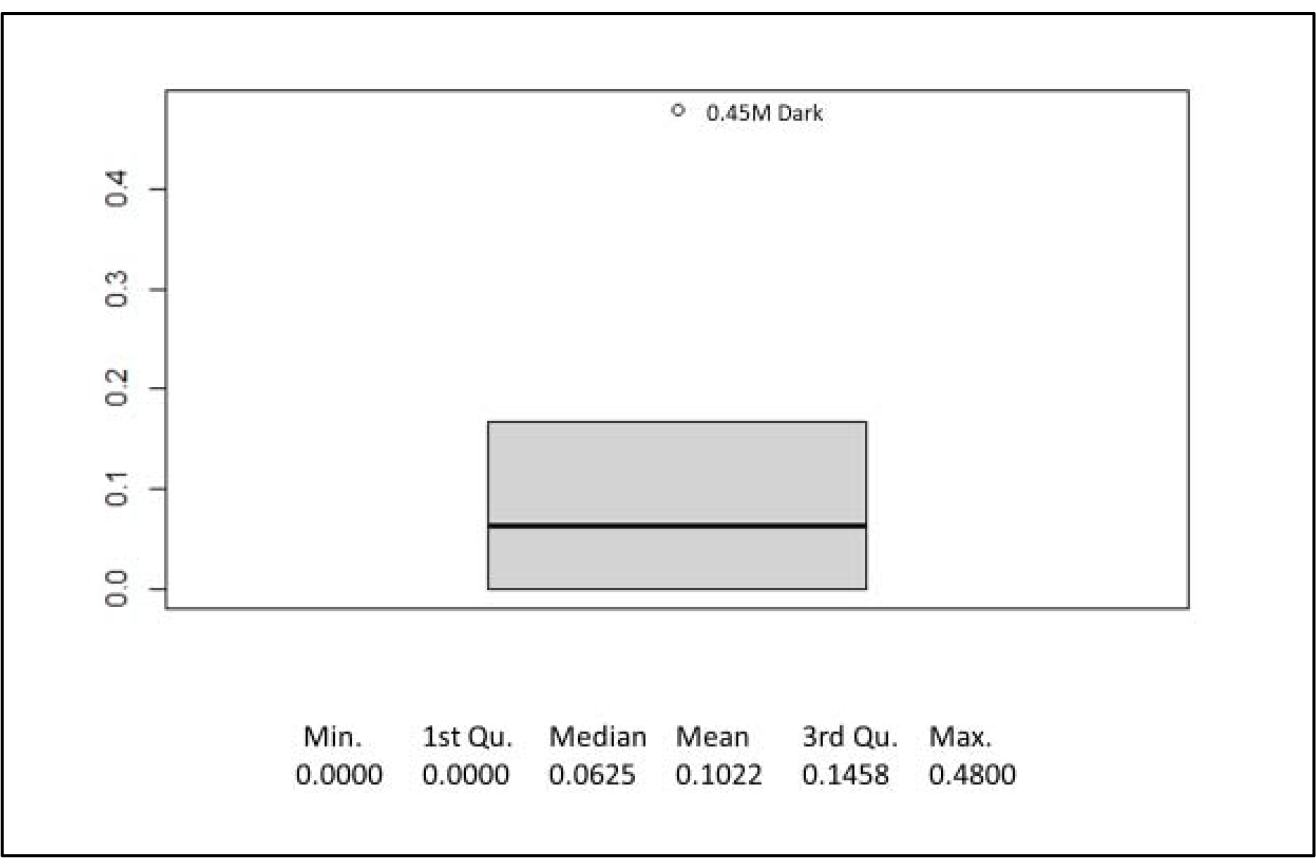
Box plot and descriptive statistics of mortality ratios of all dose groups in both dark and light experiments. The death ratio of the 0.45 M group in the dark experiment is 0.48.

**Figure S3.**
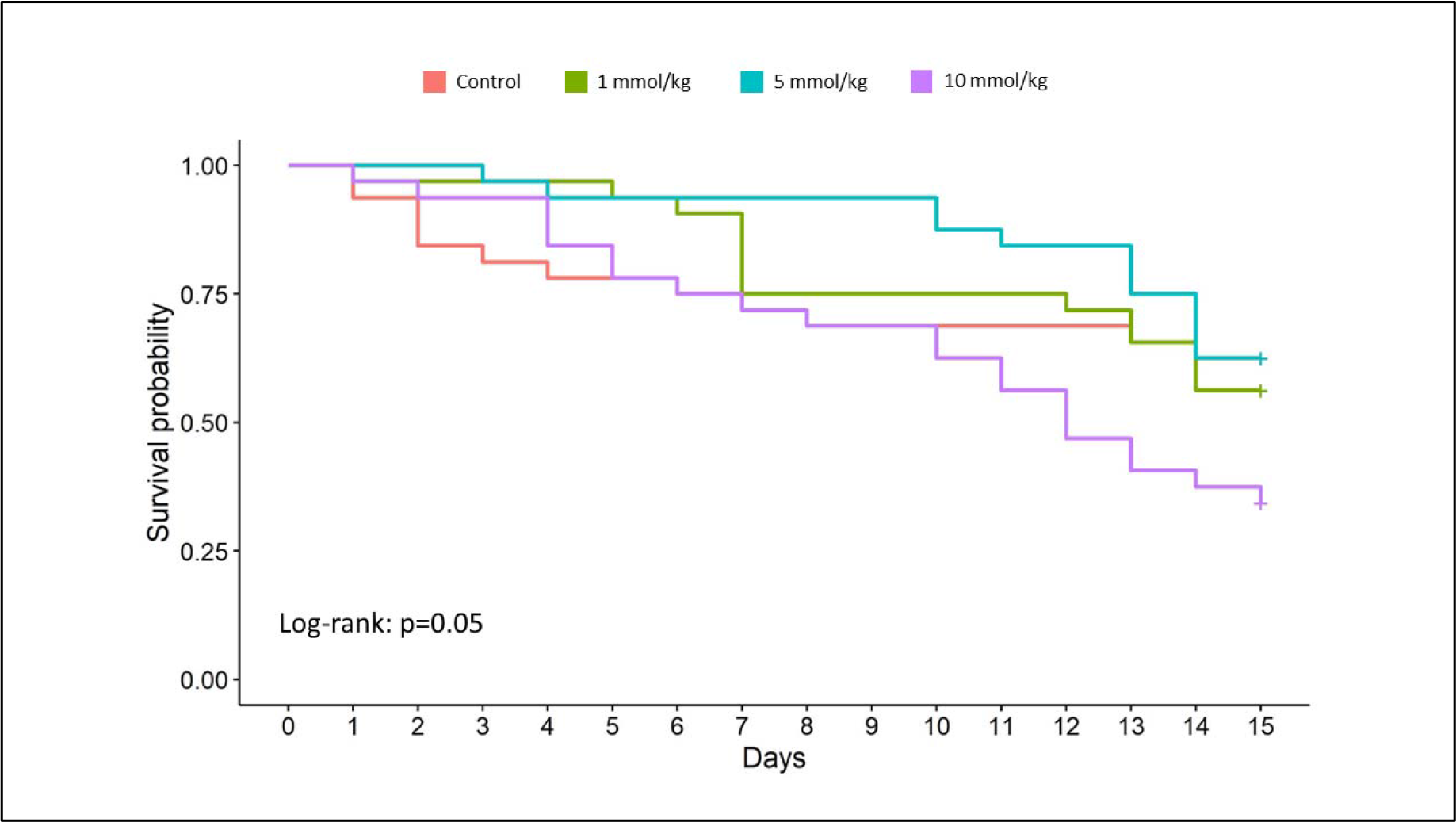
Log-rank test plot. There was no difference found between the groups (χ2 = 7.9, *p* = 0.05)

### Chronic experiments

**Table S2.**
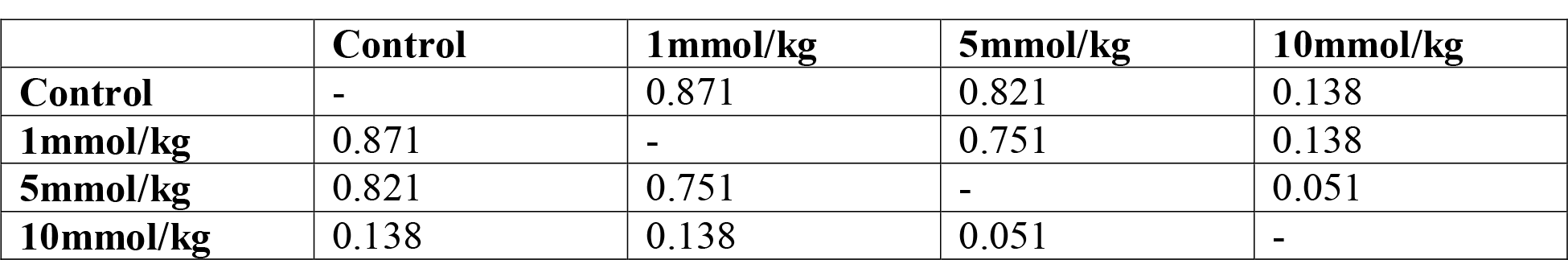
Pairwise comparisons of the survival probabilities of the doses using Log-Rank test. P values are adjusted with BH method

**Table S3.**
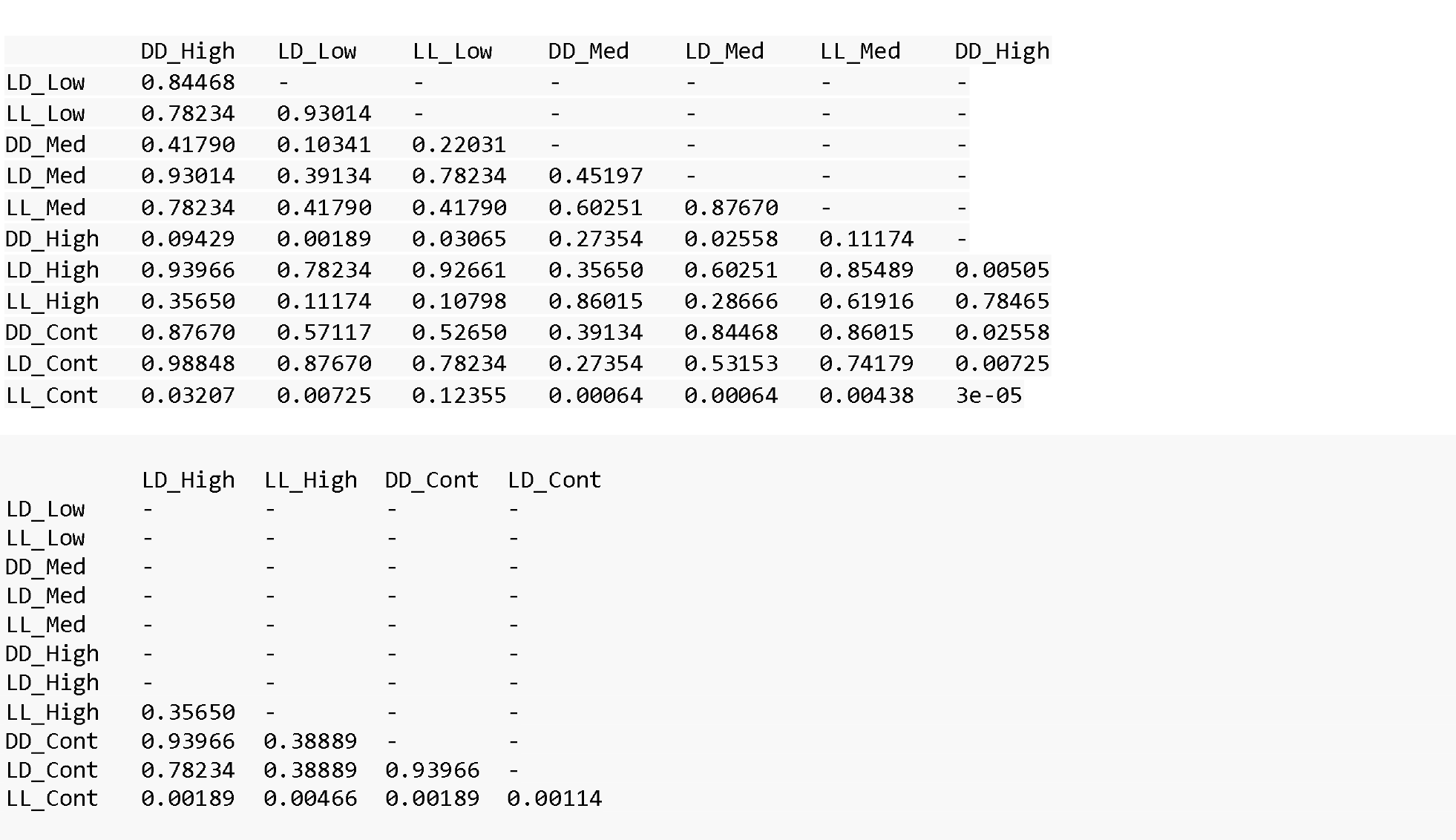
Pairwise comparisons of LMA levels in chronic experiment for all data points. The comparisons were achieved using the Wilcoxon rank sum exact test, and p values were adjusted with the Bonferroni method. In the table, High, Med., Low, and Cont. indicates 10mmol/kg, 5 mmol/kg, 1 mmol/kg, and control, respectively. LD: Light/Dark, DD: constant dark, LL constant dark.

**Figure S4.**
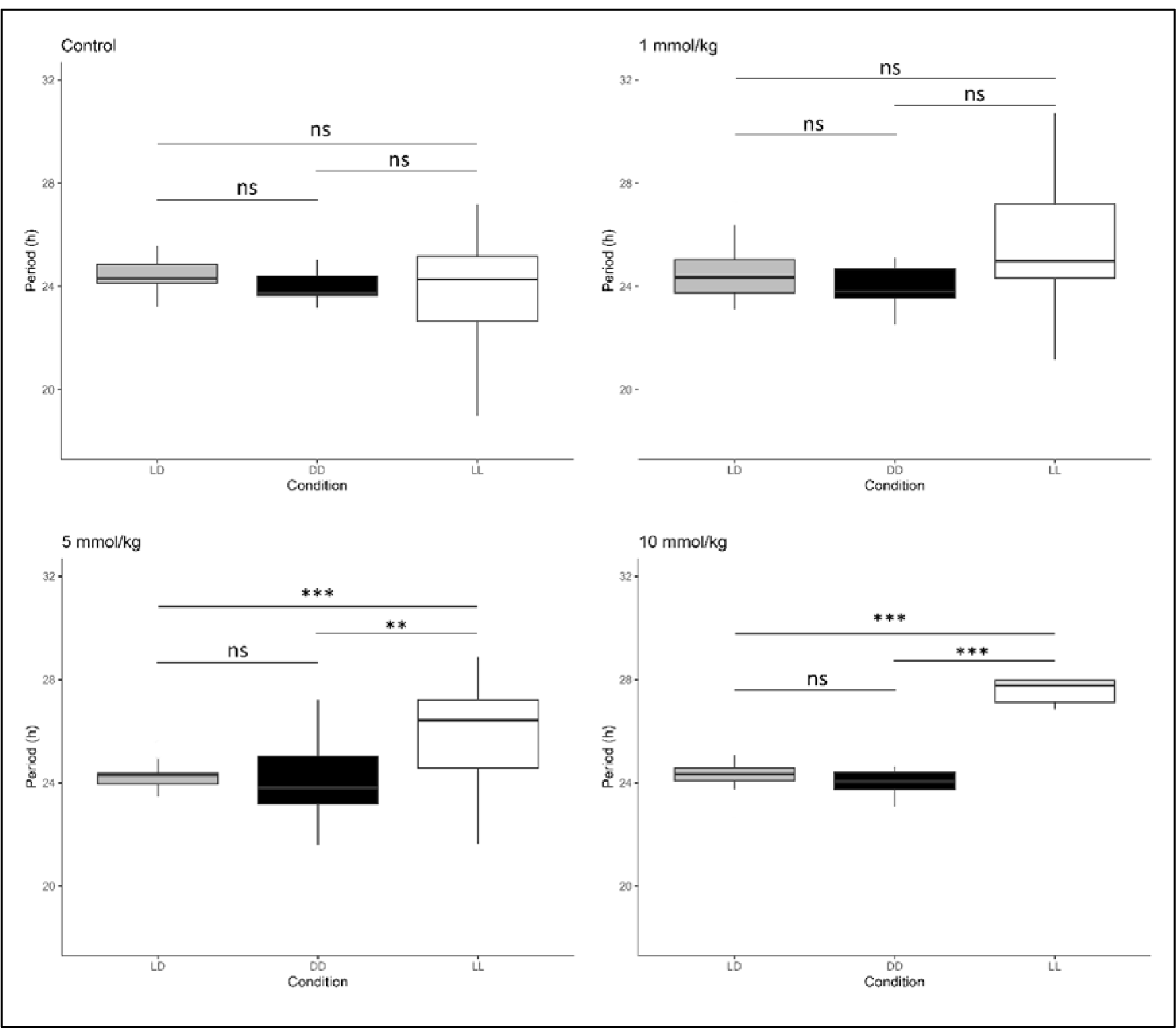
Light conditions did effective on the circadian period for medium and high dose (p < .001) but not for control and low dose groups (p > .05) according to the Kruskal-Wallis test. The length of the circadian period in LL condition was significantly higher than in other conditions according to the Dunn test, * p < .05, ** p < .01, *** p < .001, ns: not significant. Grey boxes represent LD, black boxes are DD, and whites are LL.

