## Supplemental figures and tables for "Effects of lithium on locomotor activity and circadian rhythm of honey bees"

**Acute experiments**

Table S1. Mortality ratios of the groups acute LMA experiments in the dark and light environment.

| Experiment | Treatment | Dose | Mortality Ratio |
| --- | --- | --- | --- |
| Acute LMA Experiment in Dark Environment | LiCl | 0.00M | 0.00 |
|  | LiCl | 0.05M | 0.04 |
|  | LiCl | 0.15M | 0.08 |
|  | LiCl | 0.45M | 0.48 |
|  | NaCl | 0.45M | 0.00 |
| Acute LMA Experiment in Light Environment | LiCl | 0.00M | 0.08 |
|  | LiCl | 0.05M | 0.00 |
|  | LiCl | 0.15M | 0.17 |
|  | LiCl | 0.45M | 0.17 |
|  | NaCl | 0.45M | 0.00 |

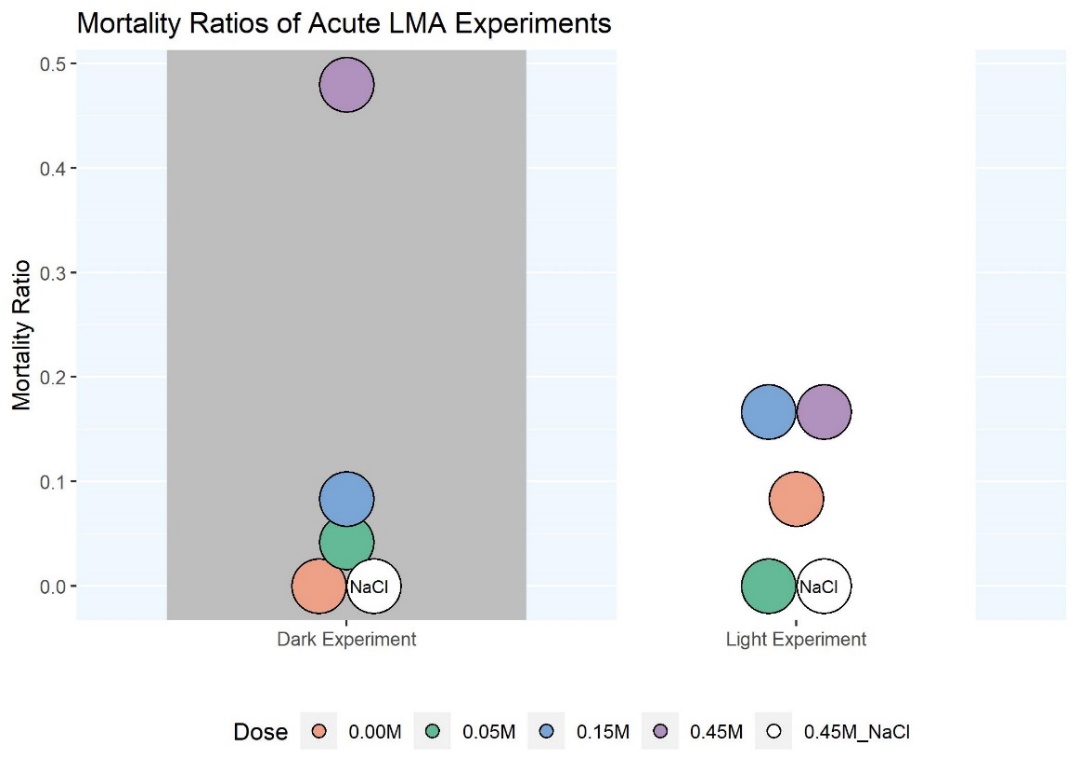

Figure S1. Mortality ratios of the groups acute LMA experiments in the dark and light environment.

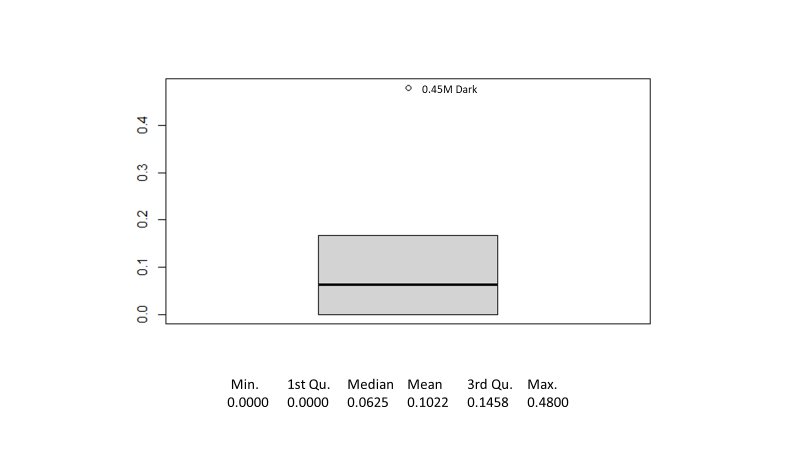

Figure S2. Box plot and descriptive statistics of mortality ratios of all dose groups in both dark and light experiments. The death ratio of the 0.45 M group in the dark experiment is 0.48.

**Chronic experiments**

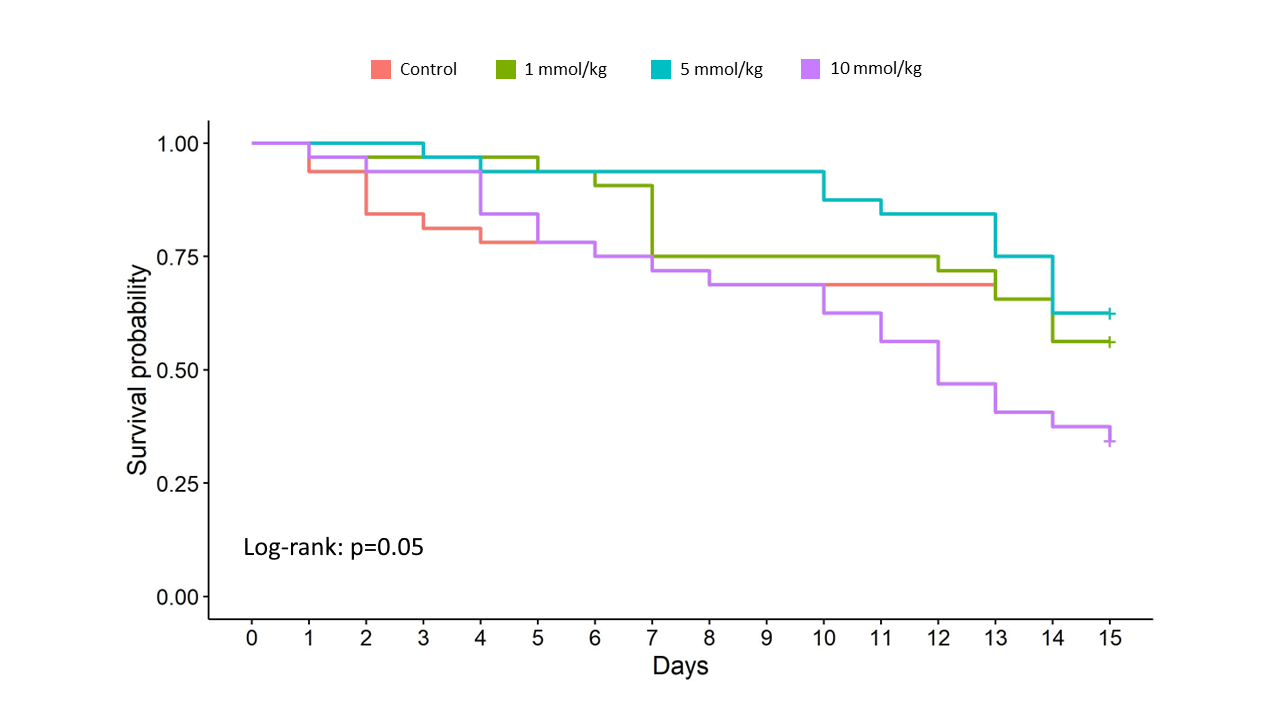

Figure S3. Log-rank test plot. There was no difference found between the groups (χ2 = 7.9, p = 0.05)

Table S2. Pairwise comparisons of the survival probabilities of the doses using Log-Rank test. P values are adjusted with BH method

|  | **Control** | **1mmol/kg** | **5mmol/kg** | **10mmol/kg** |
| --- | --- | --- | --- | --- |
| **Control** | - | 0.871 | 0.821 | 0.138 |
| **1mmol/kg** | 0.871 | - | 0.751 | 0.138 |
| **5mmol/kg** | 0.821 | 0.751 | - | 0.051 |
| **10mmol/kg** | 0.138 | 0.138 | 0.051 | - |

Table S3. Pairwise comparisons of LMA levels in chronic experiment for all data points. The comparisons were achieved using the Wilcoxon rank sum exact test, and p values were adjusted with the Bonferroni method. In the table, High, Med., Low, and Cont. indicates 10mmol/kg, 5 mmol/kg, 1 mmol/kg, and control, respectively. LD: Light/Dark, DD: constant dark, LL constant dark.

 DD_High	LD_Low	LL_Low	DD_Med	LD_Med	LL_Med	DD_High
LD_Low	0.84468	-	-	-	-	-	-
LL_Low	0.78234	0.93014	-	-	-	-	-
DD_Med	0.41790	0.10341	0.22031	-	-	-	-
LD_Med	0.93014	0.39134	0.78234	0.45197	-	-	-
LL_Med	0.78234	0.41790	0.41790	0.60251	0.87670	-	-
DD_High	0.09429	0.00189	0.03065	0.27354	0.02558	0.11174	-
LD_High	0.93966	0.78234	0.92661	0.35650	0.60251	0.85489	0.00505
LL_High	0.35650	0.11174	0.10798	0.86015	0.28666	0.61916	0.78465
DD_Cont	0.87670	0.57117	0.52650	0.39134	0.84468	0.86015	0.02558
LD_Cont	0.98848	0.87670	0.78234	0.27354	0.53153	0.74179	0.00725
LL_Cont	0.03207	0.00725	0.12355	0.00064	0.00064	0.00438	3e-05

```

          LD_High  LL_High  DD_Cont  LD_Cont
LD_Low    -        -        -        -
LL_Low    -        -        -        -
DD_Med    -        -        -        -
LD_Med    -        -        -        -
LL_Med    -        -        -        -
DD_High   -        -        -        -
LD_High   -        -        -        -
LL_High   0.35650  -        -        -
DD_Cont   0.93966  0.38889  -        -
LD_Cont   0.78234  0.38889  0.93966  -
LL_Cont   0.00189  0.00466  0.00189  0.00114

```

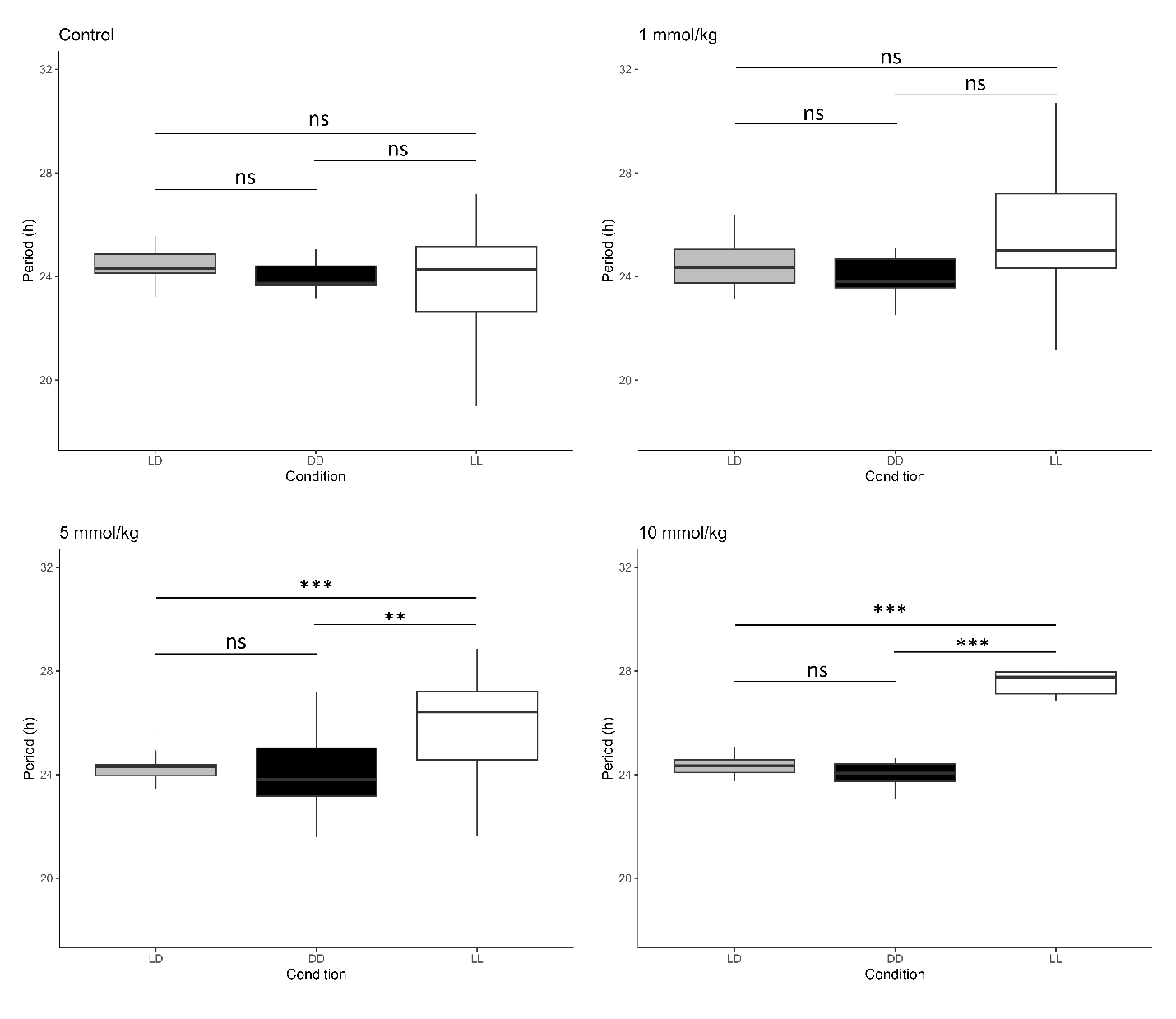

Figure S4. Light conditions did effective on the circadian period for medium and high dose (p < .001) but not for control and low dose groups (p > .05) according to the Kruskal-Wallis test. The length of the circadian period in LL condition was significantly higher than in other conditions according to the Dunn test, * p < .05, ** p < .01, *** p < .001, ns: not significant. Grey boxes represent LD, black boxes are DD, and whites are LL.
